# Mechanism and cellular function of direct membrane binding by the ESCRT and ERES-associated Ca^2+^-sensor ALG-2

**DOI:** 10.1101/2023.10.17.562764

**Authors:** Sankalp Shukla, Wei Chen, Shanlin Rao, Serim Yang, Chenxi Ou, Kevin P. Larsen, Gerhard Hummer, Phyllis I. Hanson, James H. Hurley

## Abstract

Apoptosis Linked Gene-2 (ALG-2) is a multifunctional intracellular Ca^2+^ sensor and the archetypal member of the penta-EF hand protein family. ALG-2 functions in the repair of damage to both the plasma and lysosome membranes and in COPII-dependent budding at <u>e</u>ndoplasmic <u>r</u>eticulum <u>e</u>xit <u>s</u>ites (ERES). In the presence of Ca^2+^, ALG-2 binds to ESCRT-I and ALIX in membrane repair and to SEC31A at ERES. ALG-2 also binds directly to acidic membranes in the presence of Ca^2+^ by a combination of electrostatic and hydrophobic interactions. By combining GUV-based experiments and molecular dynamics simulations, we show that charge-reversed mutants of ALG-2 at these locations disrupt membrane recruitment. ALG-2 membrane binding mutants have reduced or abrogated ERES localization in response to Thapsigargin-induced Ca^2+^ release but still localize to lysosomes following lysosomal Ca^2+^ release. *In vitro* reconstitution shows that the ALG-2 membrane-binding defect can be rescued by binding to ESCRT-I. These data thus reveal the nature of direct Ca^2+^-dependent membrane binding and its interplay with Ca^2+^-dependent protein binding in the cellular functions of ALG-2.

## Introduction

Apoptosis-linked gene (ALG-2) plays a central role in regulating various cellular processes, including plasma membrane and lysosome repair and vesicle budding at ER exit sites (ERES) (1). In the <u>E</u>ndosomal <u>S</u>orting <u>C</u>omplex <u>R</u>equired for <u>T</u>ransport (ESCRT) pathway (2, 3), ALG-2 senses Ca^2+^ released into the cytosol by plasma (4) and lysosome membrane damage (5), triggering recruitment of repair factors. ALG-2 also mediate effects of Ca^2+^ released by TRPML1 at lysosomes (6). At ERES, ALG-2 works with another penta-EF hand protein, PEF1, to regulate CUL3 and KLHL12-dependent ubiquitylation of COPII (7). Interest in the lysosome repair pathway, in particular, has intensified following recent discoveries of the role of cholesterol and ER-lysosome contact sites in repair (8, 9). Moreover, lysosome repair by the ESCRT machinery has been implicated in preventing the transmission of prion-like tau aggregates involved in Alzheimer’s disease (10).

ALG-2 consists of five repeated EF-hand motifs and belongs to the penta-EF-hand (PEF) family (1). ALG-2 contains eight α-helices and forms dimers through the pairing of EF5 (11). The binding of ALG-2 to Ca^2+^ induces conformational changes that facilitate its interaction with proline-rich segments of its targets, of which the best known are the ESCRT proteins ALIX and TSG101 and the COPII subunit SEC31A (12–14). ESCRT-I and ALIX share the same binding site (15), while SEC31A binds at a distinct site (14).

Despite that all of the above-mentioned protein-protein interactions occur in the context of membranes, and many other families of Ca^2+^-binding proteins interact directly with membranes (16–19), there has been little investigation of the role of the membrane itself in the recruitment functions of ALG-2. In the course of reconstituting Ca^2+^– and ALG-2-dependent recruitment of ESCRTs to membranes (20), we noticed that ALG-2 had an intrinsic ability to bind acidic membranes in a Ca^2+^-dependent manner even in the absence of ESCRTs. Building on this work, we investigated the mechanistic determinants and cellular function of this membrane binding activity, and whether it had any physiological role.

To address these questions, we employed a three-pronged approach. Firstly, we utilized a giant unilamellar vesicle (GUV) reconstitution system, which allowed us to leverage fluorescence microscopy in a precisely defined experimental setup. By designing ALG-2 mutants based on the X-ray crystal structure of ALG-2, we aimed to perturb its interaction with negatively charged membranes. Subsequently, molecular dynamics simulations were employed to more precisely identify the interactions responsible for the binding of ALG-2 to negatively charged membranes. Lastly, through live cell imaging techniques, we monitored the localization of ALG-2 membrane-binding mutants to ER-exit sites and lysosomes in response to Ca^2+^release.

## RESULTS

### ALG-2 mutants abrogate binding to negatively charged membranes

Based on crystal structures of ALG-2 (11, 14, 15), we predicted two regions might contribute to the membrane binding of ALG-2 to negatively charged membranes. The first is a basic patch consisting of Arg34, Lys37, and Arg39, which are exposed such that they could bind to negatively charged membranes, and the second is the exposed hydrophobic sidechain of Trp57 (Fig. 1A). We hypothesized that the electrostatic interaction of the RKR motif with the negatively charged membrane might stabilize binding of ALG-2 to acidic membranes. Once the ALG-2 is in the vicinity of the membrane, the Trp57 residue likely embeds in the membrane, reinforcing the binding of ALG-2 to negatively charged membranes.

**Figure 1.**
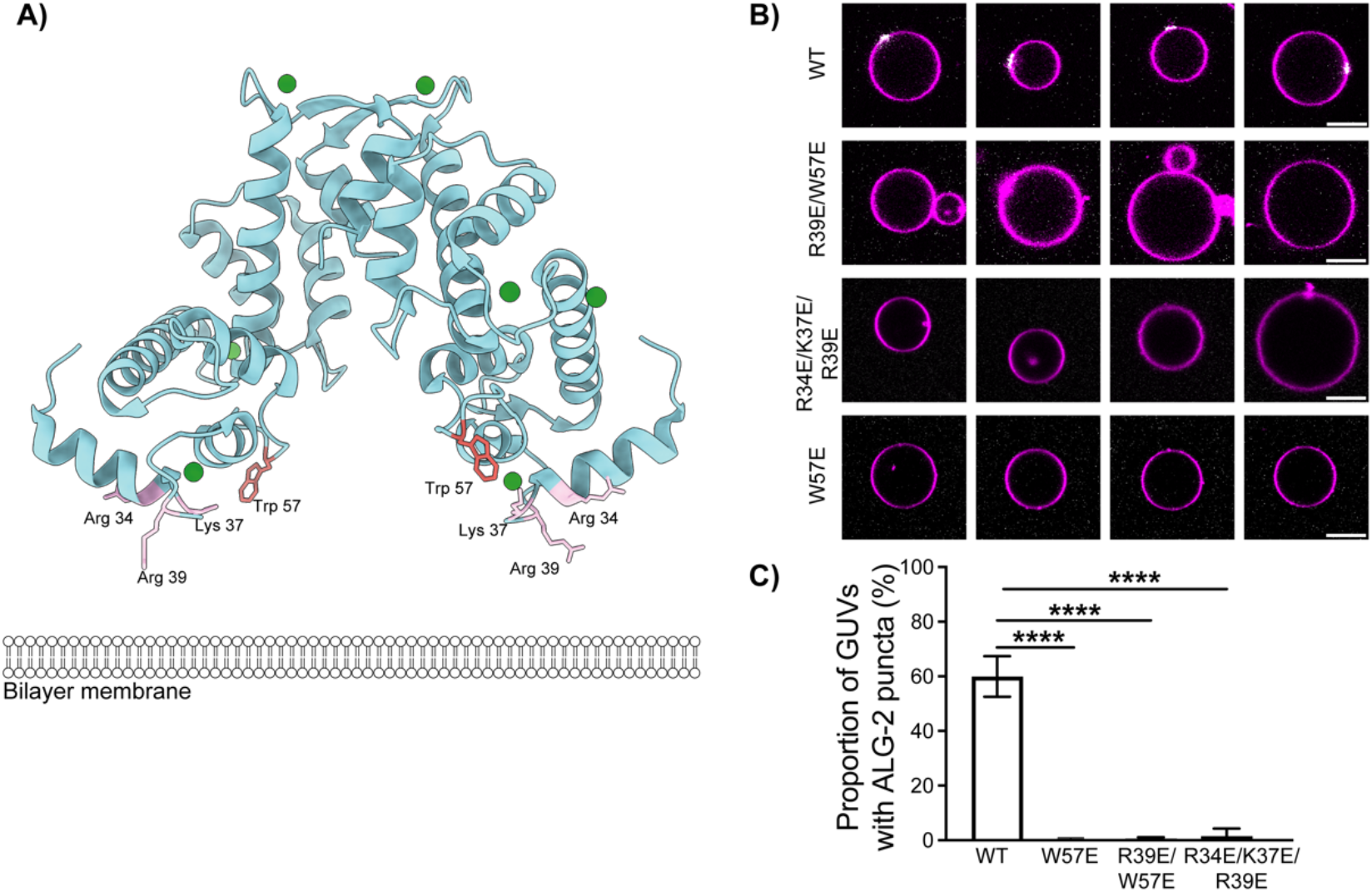
ALG-2 mutants abrogate membrane binding to negatively charged membranes in a GUV-based assay. A) Structure of Ca2+-bound ALG-2 (PDB ID: 2ZN9) with residues R39 and W57 protruding towards a cartoon bilayer membrane. B) ALG-2 A78C was fluorescently labeled with Atto 488 at the introduced Cys78. All mutants shown here were made in the A78C background. Wild-type and the indicated mutant ALG-2 proteins were incubated with 30% DOPS containing GUVs. C) The proportion of GUVs that had at least one ALG-2 punctum (white) on their periphery (magenta) were plotted for fluorescently labeled WT ALG-2 (n = 1388 GUVs), W57E (n = 395 GUVs), R34E/K37E/R39E ALG-2 (n = 741 GUVs), and R39E/W57E ALG-2 (n = 1135). The data are shown as mean ± SD (vertical line). All results are from at least three independent experiments. P ≤ 0.0001 (****). The scale bar is 10 μm.

GUV membrane binding was tested for the point mutant W57E, the triple mutant R34E/K37E/R39E, and the double mutant R39E/W57E. All of these mutants were made in the background of an introduced Cys78 for fluorophore labeling at a location distal to the putative membrane binding face. Proteins were expressed at the same levels and behaved essentially the same in purification, consistent with a lack of gross changes in protein stability. The AlphaFold2 (AF2) (21) predicted model of R34E/K37E/R39E ALG-2 superimposed upon the crystal structure of des3-20 WT ALG-2 [Protein Data Bank (PDB – 2ZN9) resulted in a root mean square deviation (RMSD) of 0.7 Å across the ordered backbone atoms from residues 24 to 188 (Fig. S1). The lack of structural perturbation was expected and is consistent with the exterior location of the mutated side-chains. We incubated GUVs containing 30% 1,2-dioleoyl– sn-glycero-3-phospho-L-serine (DOPS) along with 69.5% 1,2-dioleoyl-sn-glycero-3-phosphocholine (DOPC) and 0.5% Atto 647N dye-labeled 1,2-dioleoyl-sn-glycerol-3-phosphoethanolamine (DOPE) with WT ALG-2 and all its mutants (Atto 488; 200 nM) for 15 min in reaction buffer. On imaging, the WT ALG-2 formed puncta on the membrane as reported previously (20). The triple charge reversal mutant ALG-2^R34E/K37E/R39E^ abrogated the membrane binding of ALG-2.

Moreover, the double mutant, R39E/W57E ALG-2 with a mutation in the basic patch and of the hydrophobic residue and W57E ALG-2 single mutation of the hydrophobic residue were also sufficient to abrogate the binding of ALG-2 to the negatively charged membranes (Fig. 1B, C). Previously, the N-terminal region of ALG-2 rich in alanine, glycine, and proline residues has been speculated to contribute to the membrane binding of ALG-2 (22). To test this, we generated the ΔN23 mutant of ALG-2 and did the same experiment with GUVs. We found that the ΔN23 mutant of ALG-2 still binds the negatively charged membranes similar to that of WT ALG-2 (Fig. S2). Taken together, this points to a concerted membrane binding effort of ALG-2, whereby electrostatic interactions position the ALG-2 dimer close to the membrane, and then the insertion of the tryptophan residue in the hydrophobic core of the bilayer membrane cements the ALG-2 dimer to the membrane.

### A triad of basic residues in EF1 drives electrostatic interactions with membrane

In all-atom molecular dynamics (MD) simulations, Ca^2+^-bound ALG-2 was observed to bind readily to negatively charged membranes consisting of 30% DOPS and 70% DOPC. A DOPC only control for the Ca^2+^-bound (holo) WT ALG-2 showed a statistically significant reduction in contacts compared to the 30% DOPS and 70% DOPS membranes (Fig. S3). This confirms our previous observation about the role of negative charge density in driving the calcium dependent membrane recruitment of ALG-2 (20).

In the MD simulations, stable interactions were established mainly through EF1 (and to a lesser extent EF3) from each subunit of the dimer (Fig. 2A and Fig. S4A). The three basic residues forming the RKR motif in EF1, namely Arg34, Lys37, and Arg39 examined above (Fig. 1A) initiated membrane association of ALG-2 through direct contacts with membrane lipids (Fig. 2A). Ca^2+^ ions bound to EF1 and EF3, partially encaged by their coordinating residues, were typically held at a distance of > 0.5 nm from the nearest lipid atom (Fig. S4B). Direct bridging of membrane lipids by Ca^2+^ did occur, albeit infrequently, with a total of six instances observed across six 2 µs simulation replicates. It is not possible, however, to rule out that more extensive sampling might reveal a more kinetically stable coordination complex. In each case, one or two phosphate oxygen atoms from a PS molecule replaced the Ca^2+^–coordinating Asp/Glu oxygen(s) of the EF hand (Fig. S4C). These direct coordination events (applying a distance cut-off of 0.35 nm) were generally short-lived (< 2 ns in duration), with one longer interaction persisting for ∼350 ns. No direct Ca^2+^ coordination occurred for PC lipids present in the membrane. Compared to the more active role of Ca^2+^ in mediating lipid interactions of C2 domain proteins (23, 24) and annexins (16), our simulation results suggest a mechanism of Ca^2+^-dependent membrane binding principally through an electrostatic effect. The binding of Ca^2+^ ion switches the sign of the surface electrostatic potential around the EF hands from strongly negative to positive (Fig. 2B). Drawn to the negatively charged membrane (Fig. S4D), holo ALG-2 then forms tight interactions with lipids via the RKR motif and a flanking aromatic residue. As might be expected, all three residues of the RKR motif show a strong preference for the negatively charged PS over PC (Fig. 2C). Among other residues that make frequent membrane contacts (including Asn30, Asn54, and Trp57), we observed the aromatic side chain of Trp57, in particular, inserted between lipid headgroups and interacted with the acyl core of the membrane (Fig. 2A and D). These hydrophobic interactions of Trp57 likely help to anchor ALG-2 to the target membrane.

**Figure 2.**
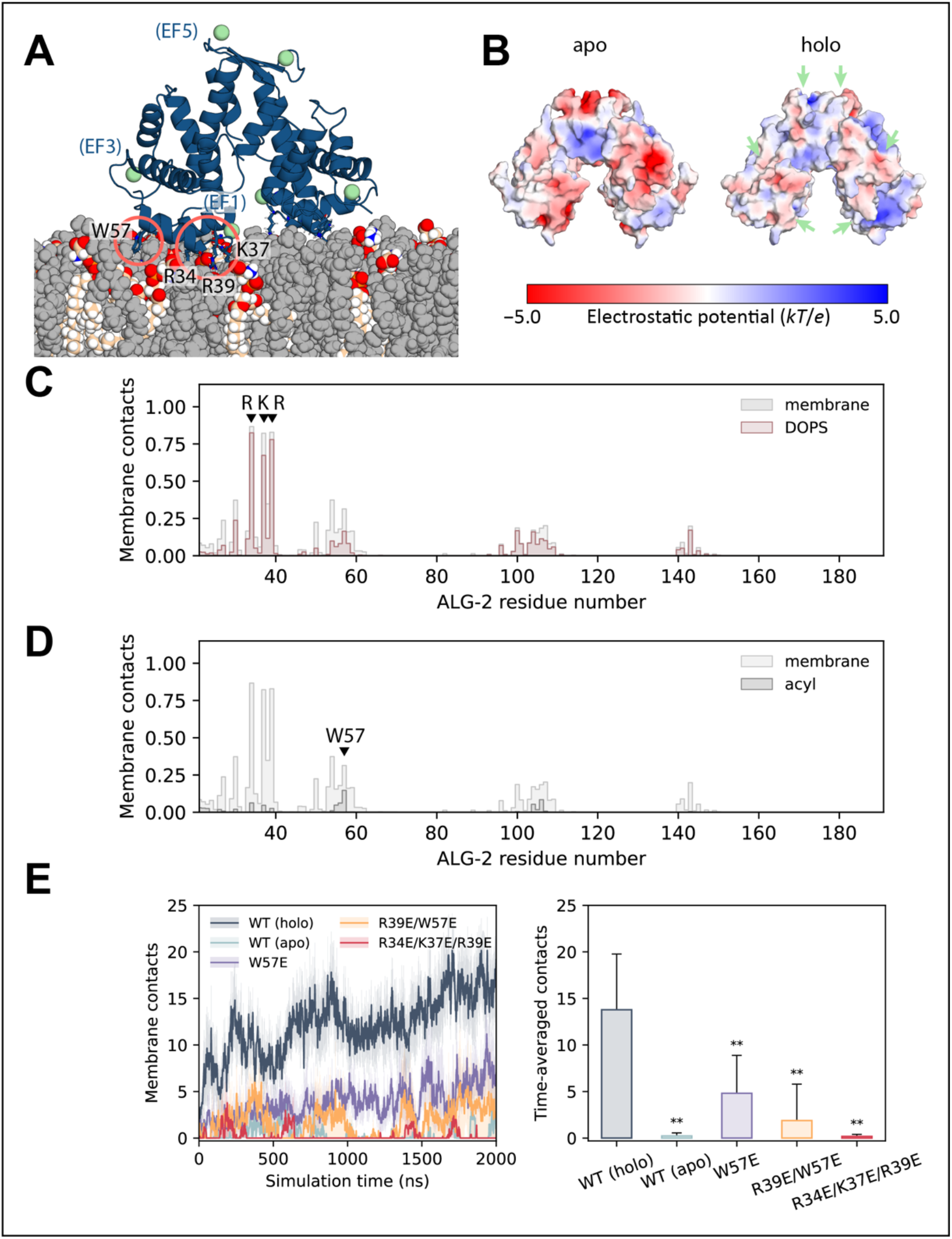
Molecular dynamics simulations of ALG-2 membrane interactions. (A) Ribbon representation of Ca^2+^-bound ALG-2 interacting with membrane lipids (spheres) upon spontaneous membrane binding during molecular dynamics simulation. PS lipids are colored by atom type, with PC lipids in gray. Key membrane-interacting residues are highlighted. (B) Surface electrostatic potential of apo and Ca^2+^-bound (holo) ALG-2. Positions of bound Ca^2+^ ions are indicated with green arrows. (C) Mean frequency of membrane and DOPS lipid contacts by each ALG-2 residue along the amino acid sequence, calculated as a percentage of analyzed frames from the final 1 µs of each of six 2 µs simulations, sampling at 2-ns intervals. (D) Mean frequency of membrane lipid and acyl tail contacts by each ALG-2 residue along the amino acid sequence. (E) Number of ALG-2 residues forming membrane contacts over the course of simulation replicates, comparing between the apo and Ca^2+^-bound (holo) forms of wild-type (WT) ALG-2 and with mutants (holo). The mean (solid lines) and standard errors (semi-transparent shading) are plotted over time for six simulation replicates of WT ALG-2 and five of each mutant. Time-averaged membrane contacts and standard deviations are also calculated across the final 1 µs of 2 µs simulation replicates of each structure. Statistically significant (0.001 < *p* < 0.01) reductions in membrane binding compared with the Ca^2+^-bound WT protein, as assessed by one-tailed Student’s t-tests, are denoted by asterisks (**).

### Mutations reduce ALG-2 membrane binding in molecular dynamics simulations

In agreement with results from GUV experiments, the mutations W57E, R39E/W57E, and R34E/K37E/R39E in and near the RKR motif yielded dramatically reduced membrane recruitment in all-atom MD simulations. The R34E/K37E/R39E triple mutant, in particular, shows only transient membrane interactions at a level comparable to that of the apo protein (Fig. 2E).

### ALG-2 mutants do not concentrate on ER exit sites following calcium efflux from the ER

ALG-2 is known to be recruited to ER exit site (ERES) in response to local Ca^2+^ release (1, 13). ALG-2 binds COPII protein Sec31A at ERES and regulates ER to Golgi vesicle transport (25, 26). However, how ALG-2 associates with the ERES membrane is still unclear. To test whether requirements for ALG-2 binding to ERES parallel those of ALG-2 binding to liposomes, we expressed mCherry-ALG-2 WT or W57E in HeLa cells and examined responses to Thapsigargin (TG) treatment. TG induces passive ER Ca^2+^ release by inhibiting uptake into the ER via SERCA Ca^2+^-ATPases and is known to trigger ALG-2 recruitment to ERES. As expected, ALG-2 WT formed puncta in response to TG that co-localized with Sec31A (Fig. 3A). In contrast, W57E mutant ALG-2 did not respond (Fig. 3A). To follow the dynamics of this ALG-2 recruitment, we used live-cell imaging and observed the expected rapid accumulation of WT but not W57E mCherry-ALG-2 puncta following TG treatment (Fig. 3B&C). We also examined the response of R34E/K37E/R39E ALG-2 to TG and found that its response was reduced compared to that of WT (Fig 3D&E). These data show that all of the mutants that abrogate binding to GUVs in vitro also abrogate or at least reduce localization to ERES in response to ER Ca^2+^ release.

**Figure 3.**
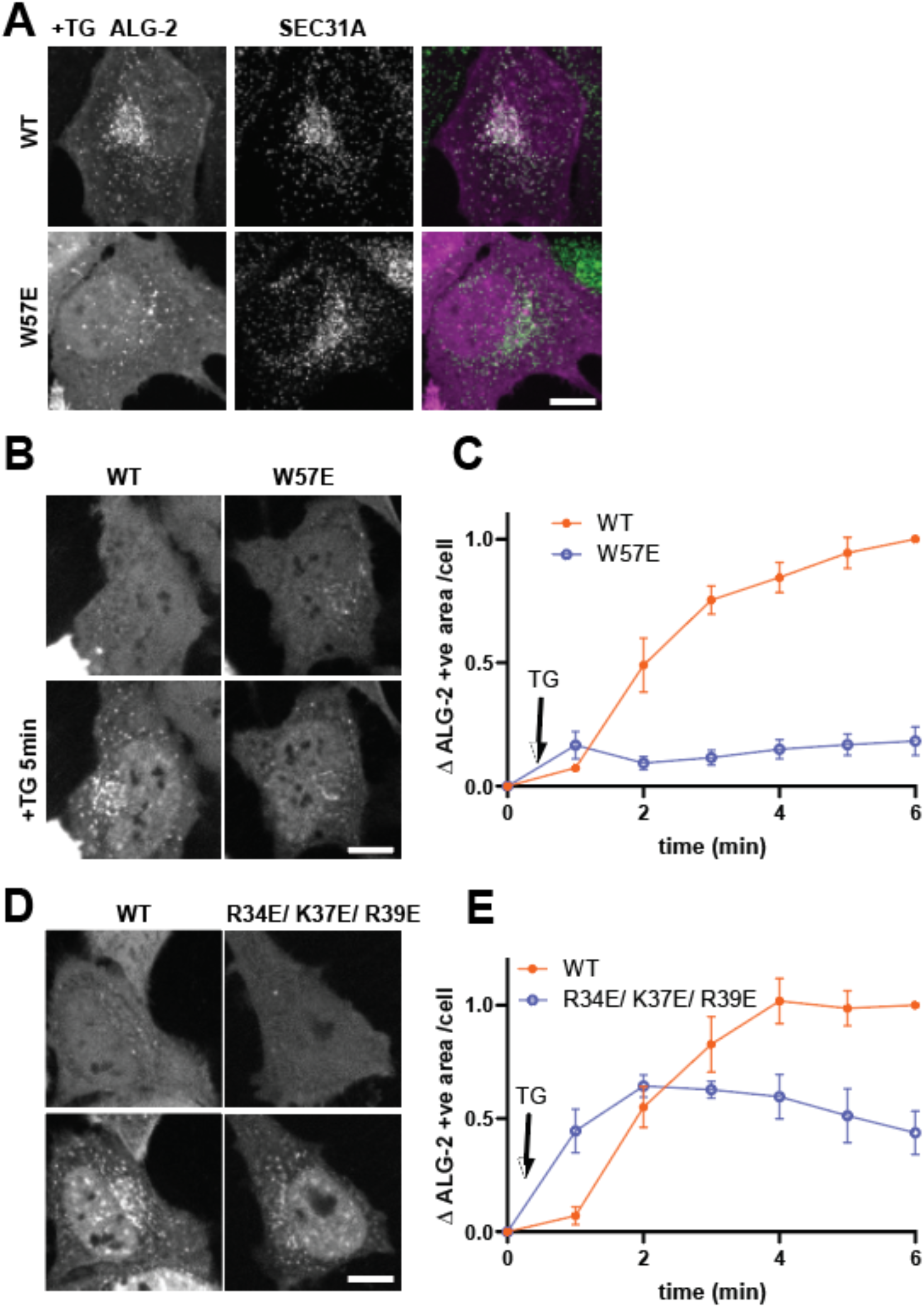
ALG-2 mutants do not concentrate at ER exit sites following Thapsigargin (TG)-induced calcium release. (A) HeLa cells expressing mCherry-ALG-2 WT or W57E were treated with TG and stained for SEC31A. (B – E) Live imaging of HeLa cells expressing mCherry-ALG-2 WT, W57E or R34E/ K37E/ R39E were imaged at 1 min intervals. (B) and (D) show cells before and after 5 min of TG treatment. (C) and (E) show mCherry-ALG-2 signal above threshold (mean ± SEM). 38 cells (WT) and 58 cells (W57E) from 3 experiments were analyzed in (C). 49 cells (WT) and 60 cells (R34E/ K37E/ R39E) from 3 experiments were analyzed in (E). Scale bar is 10 μm.

### ALG-2 mutants respond normally to lysosomal Ca^2+^ release

Locally triggered Ca^2+^ release is also reported to recruit ALG-2 to lysosomes where it further engages ESCRT proteins (5,20). To investigate the effect of the ALG-2 membrane binding mutations on its response to lysosomal Ca^2+^ release, we again performed live-cell imaging experiments. We transfected either mCherry-ALG-2 WT or ALG-2 mutants along with fluorescently tagged ESCRT-III protein IST1 into ALG-2 KO HeLa cells. We then used a TRPML1 agonist, ML-SA5 (27), along with TG and observed the expected recruitment of WT ALG-2 together with IST1 (Fig. 4A) and in the absence of IST1 (Fig. S5). All ALG-2 mutants were similarly recruited along with IST1 to lysosomes at a level similar to that of WT ALG-2 (Fig 4B, C&D). This suggests that recruitment of ALG-2 mutants to lysosomal membranes and their association with ESCRT proteins is not affected by the ALG-2 membrane binding mutations.

**Figure 4.**
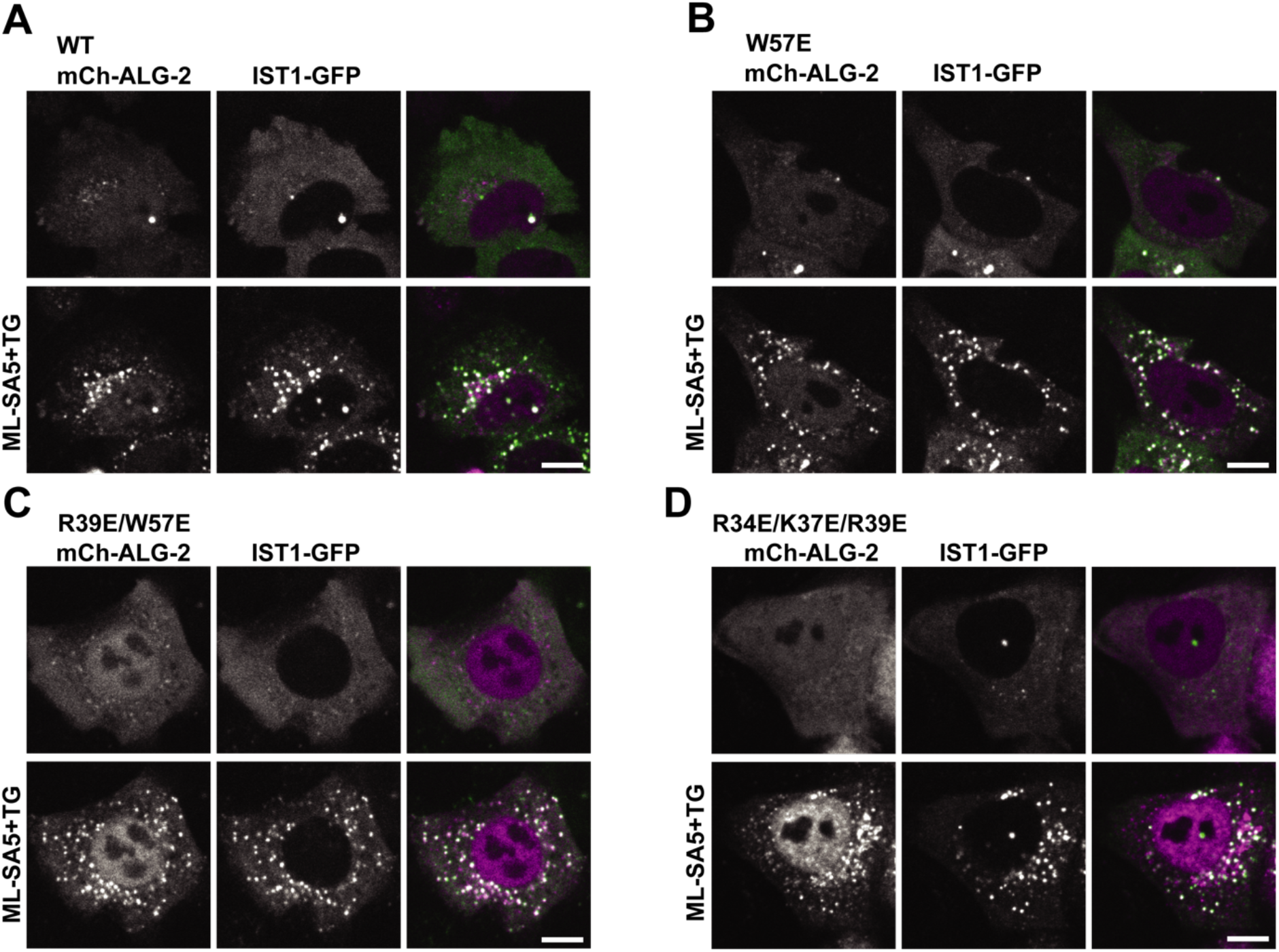
ALG-2 mutants are recruited normally to lysosomes in response to triggered Ca^2+^ release. ALG-2 KO HeLa cells were co-transfected with IST1-mGFP and mCherry-ALG-2 WT (A), W57E (B), R39E/ W57E (C) or R34E/K37E/ R39E (D). Cells were imaged live before and 15 min after incubation with ML-SA5 and TG. The scale bar is 10μm.

### ESCRT-I rescues the membrane binding ability of ALG-2 mutants

The abrogation of ERES but not lysosome recruitment of the W57E ALG-2 mutant might be explained by the presence of ESCRT machinery in the latter case. We hypothesized that the Ca^2+^ triggered recruitment of ESCRT machinery to lysosomal membrane could compensate for the decreased membrane binding of designed ALG-2 mutants.

To test compensation *in vitro*, we first incubated ESCRT-I (Cy3; 50 nM) and W57E ALG-2 (Atto 488; 200 nM) individually with 30% DOPS-containing membranes. We neither observed the recruitment of ESCRT-I nor W57E ALG-2 on the membrane individually (Fig. 5A; top and middle row, 5B) under our experimental conditions. Next, we repeated the same experiment but with W57E ALG-2 (Atto 488; 200 nM) and ESCRT-I (Cy3; 50 nM) added together on the membrane. We observed clear colocalized puncta of ESCRT-I and W57E ALG-2 on the membrane (Fig. 5A; bottom row, 5B and C). This suggests that the reduction in the membrane affinity of the W57E ALG-2 mutant is likely compensated by the ability of ESCRT-I itself to bind directly to membrane, for example, via the <u>M</u>VB12-<u>A</u>ssociated <u>β</u>-<u>p</u>rism (MABP) motif in ESCRT-I (28).

**Figure 5.**
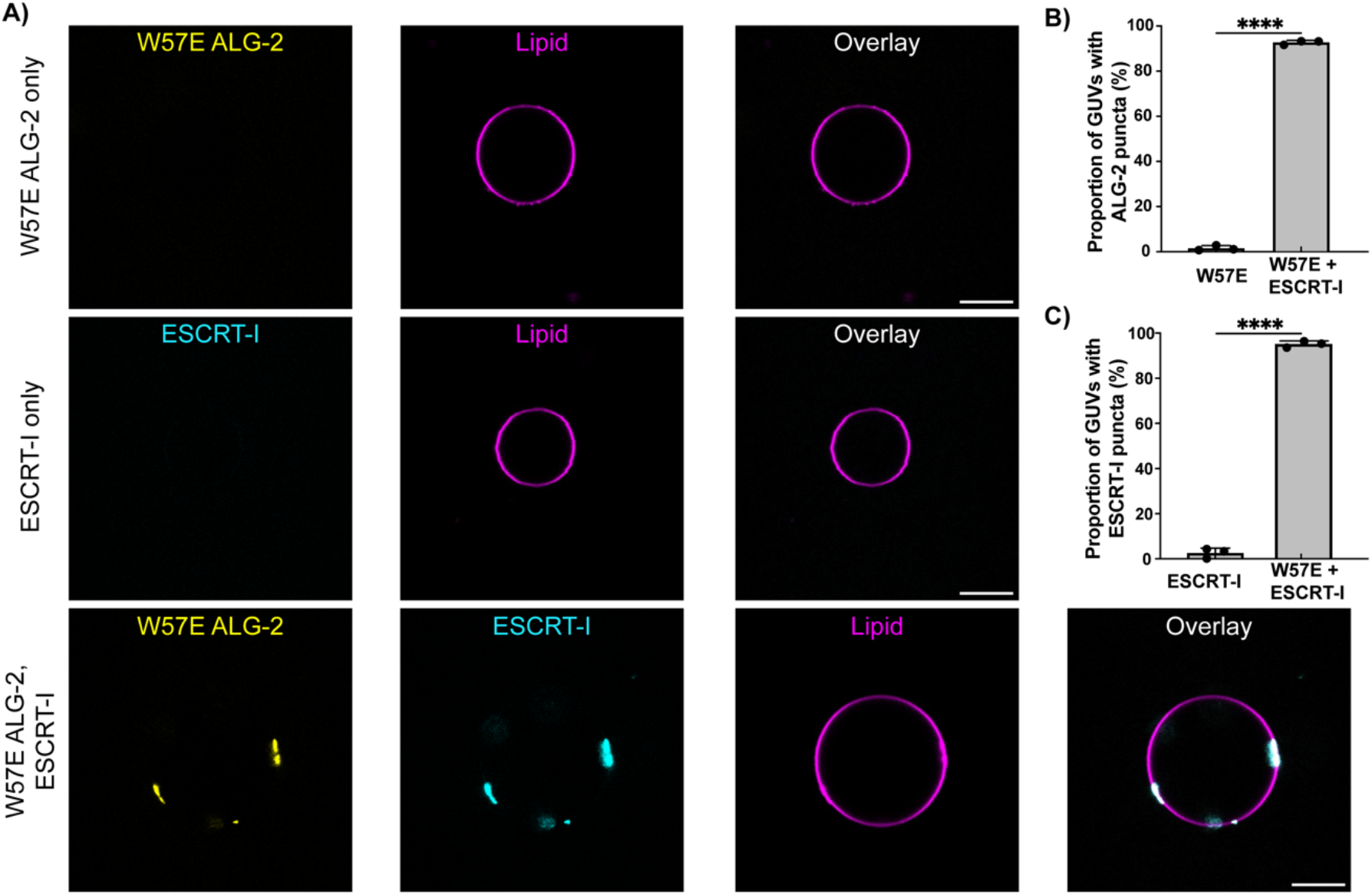
ESCRT-I rescues the membrane recruitment of ALG-2 mutants. A) (Top Row) Fluorescently labeled W57E ALG-2 (Atto 488; 200 nM) was mixed with 30% DOPS containing GUVs and imaged. No recruitment of W57E ALG-2 on the membranes was observed. (Bottom row) Upon the addition of ESCRT-I (Cy3; 50 nM) to the GUVs incubated with W57E ALG-2 (Atto 488; 200 nM), clear puncta of W57E ALG-2 and ESCRT-I onto the membranes, which also colocalize, are observed. B) The proportion of GUVs that had at least one ALG-2 punctum on their periphery were plotted for fluorescently labeled W57E ALG-2 only (n = 395 GUVs) and W57E ALG-2 with ESCRT-I (n = 332 GUVs). C) Shows the proportion of GUVs with ESCRT-I punctum on their periphery with ESCRT-I only (n = 372 GUVs) and ESCRT-I with W57E ALG-2 (n = 332 GUVs). The circles on the bar charts represent independent data points and the data are shown as mean ± SD (vertical line). All results are from at least three independent experiments. P ≤ 0.003 (***), ns = not significant. (Scale bar, 10 μm.) The scale bar is 10 μm.

## Discussion

While ALG-2 has long been understood to undergo Ca^2+^-triggered protein-protein interactions, the role of Ca^2+^ in promoting its direct binding to acidic membranes has not been appreciated until recently. Our findings show that for ALG-2, the role of Ca^2+^ binding is to neutralize charge repulsion between the electronegative apo-ALG-2 protein and acidic membranes. Both the lysosome and ER membranes are negatively charged, as is the plasma membrane (29–31). This electrostatic neutralization mechanism contrasts to the direct Ca^2+^ bridging that has been characterized for annexins and C2 domains (16–19). In the case of ALG-2, direct Ca^2+^ – phospholipid coordination was briefly observed in simulation, but only rarely and for a few ns at a time. Thus, ALG-2 defines a mechanistically novel class of Ca^2+^-stimulated membrane-binding protein.

ALG-2 is recruited both to lysosomal membrane and to ERES upon calcium efflux. A key distinction between these events is the absence ESCRT machinery at ERES. ALG-2 has three substrate binding pockets termed ALG-2 binding sites (ABS), ABS-1, –2, and –3. TSG101 and ALIX bind to pockets 1 and 2, whereas Sec31 binds to pocket 3. Trp 57 is at the edge of binding pocket 3, however, ALG-2 W57A has essentially normal binding to Sec31 *in vitro* (14). Furthermore, Arg 34, Lys 37, and Arg 39, which do have reduced ER recruitment, are located outside of pocket 3. The loss of ERES recruitment seen in these mutants can thus only reasonably be explained by the loss of membrane binding rather than a defect in Sec31 interaction.

ESCRT components, including ESCRT-I, have intrinsic membrane binding affinity. The MABP domain of the MVB12 subunit of ESCRT-I mediates its interaction with anionic lipids (28). We previously found that ESCRT-I binds to 30% DOPS-containing membranes under the current experimental conditions at concentrations of 100 nM and 200 nM, but not 50 nM (20). Therefore, it is plausible that the downstream recruitment of ESCRT machinery assists the recruitment of ALG-2 to membranes in a positive feedback loop. The rescue of the membrane-binding ability of otherwise membrane non-binding ALG-2 mutants upon the addition of ESCRT-I in our GUV-based assay is consistent with this hypothesis. This supports a primary role for Ca^2+^-stimulated ALG-2-ESCRT interaction, with a more secondary and ancillary role for Ca^2+^-stimulated membrane binding in lysosome biology.

## Materials and Methods

### Materials

The lipids 1,2-dioleoyl-sn-glycero-3-phosphocholine (DOPC) and 1,2-dioleoyl-sn-glycero-3-phospho-L-serine (sodium salt) (DOPS) were obtained from Avanti Polar Lipids (Alabaster, AL). Lipid fluorophore 1,2 – Dioleoyl-sn-glycero-3-phosphoethanolamine labeled with Atto 647 (Atto 647 DOPE) was purchased from Sigma Aldrich (St. Louis, MO). Thapsigargin was used at 2 μM (AC328570010, Thermo Scientific). HEPES, NaCl, EGTA, and fatty acid-free bovine albumin (BSA) were obtained from Fisher Scientific (Rochester, NY). All commercial reagents were used without further purification.

### GUV formation

GUVs containing DOPC (69.5 mol%), DOPS (30 mol%), and the lipid fluorophore Atto 647 DOPE (0.5 mol%) were prepared in 270 mOsm sucrose using PVA-gel hydration-based method as in Weinberger et al. (32). Briefly, lipids were mixed in chloroform at a total concentration of 1 mM. The 40 μL solution of the lipid mixture was spread on a 5% w/v PVA film dried on a 25 x 25 mm coverslip (VWR, Radnor, PA) and then put under vacuum for at least 2 h to form a dry lipid film. The dried lipid film was hydrated with 500 μL of 270 mOsm sucrose solution for 2 h at room temperature to produce GUV dispersion which was collected and stored in a 1.5 ml microcentrifuge tube.

### Protein purification

ALG-2 and all the mutants of ALG-2 (W57E; R34E/K39E/R39E; R39E/W57E; ΔN23) were purified based on the ALG-2 purification protocol described in McGourty et al. (7). Briefly, N-terminal 6x His tagged ALG-2 is expressed in *E. coli* BL21 (DE3) cells in LB medium supplemented with kanamycin (50 μg/ml), induced at 0.8 OD with 0.5 mM IPTG at 37 °C for 3 h. After lysis through tip sonication in lysis buffer (50 mM Tris pH 7.4, 150 mM NaCl, 0.2 mM TCEP) the expressed protein was extracted from the supernatant using NiNTA resin (QIAGEN, Germantown, MD). The resulting eluate from the NiNTA resin using lysis buffer supplemented with 250 mM Imidazole pH 7.4, was loaded onto the Superdex 75 16/60 column (GE Healthcare) for gel filtration. Subsequently, the resulting solution was purified using anion exchange chromatography using 5 ml HiTrap Q HP (Cytiva, Marlborough, MA). Finally, the eluate was loaded on an equilibrated Superdex 75 16/60 column (GE Healthcare), and the protein purity was assessed using sodium dodecyl sulfate-polyacrylamide gel electrophoresis (SDS-PAGE). The concentration of the purified protein was calculated by measuring the absorbance at 280 nm. Finally, the protein was concentrated at approximately 50 μM and stored at –80°C in small aliquots.

ESCRT-I (TSG101, VPS28, VPS37B, and MVB12A) was expressed in HEK293 cells. ESCRT-I had a Strep-tagged VPS28 subunit and was initially purified on StrepTactin Sepharose (IBA), followed by gel filtration chromatography on a Superdex 200 16/60 column (GE Healthcare), in 50 mM Tris pH 7.4, 300 mM NaCl, 0.1 mM TCEP.

Fluorophore labeling was performed overnight at 4°C using cysteine reactive dyes on engineered cysteines (A78C for ALG-2 and all its mutants) or native surface-exposed cysteines (ESCRT-I). Specifically, Atto 488 maleimide (Sigma-Aldrich) was used for labeling ALG-2 (and all its mutants) and sulfo-Cy3 maleimide (Fischer Scientific, Hampton, NH) for labeling ESCRT-I. Excess dye was removed by sequentially passing the protein-dye mixture through two PD10 columns (Cytiva, Marlborough, MA). The final step in every protein purification was gel filtration chromatography to ensure the monodisperse state of the (labeled) protein. Labeling efficiencies were normally around 25% for ALG-2 and all its mutants and >75% for ESCRT-I.

### Reconstitution Reactions and Confocal Microscopy

The incubation reactions were set up in a microcentrifuge tube at room temperature before transferring to a Lab-Tek II chambered cover glass (Fisher Scientific) for imaging. The imaging chamber was pre-coated with a 5 mg/ml solution of fatty acid-free Bovine Serum Albumin for 30 min and washed three times with the reaction buffer (25 mM HEPES at pH 7.4, 125 mM NaCl, and 0.2 mM TCEP, 280 mOsm) before transferring the reactants from the microcentrifuge tubes. 15 μL of GUVs were mixed with 120 μL of reaction buffer containing proteins at concentrations stated in the results section. After 15-minute incubation images were acquired on a Nikon A1 confocal microscope with a 63× Plan Apochromat 1.4 numerical aperture (NA) objective. Three replicates were performed for each experimental condition in different imaging chambers. Identical laser power and gain settings were used for each set of replicates. The reconstitution experiments were performed within 48 h of GUV preparation to limit the photooxidation of lipids. All experiments were performed at room temperature.

The images for cell experiments were acquired on an Olympus IX83 microscope with a Yokogawa CSU-W1 spinning disk confocal scanner unit, an Olympus PlanApo 60x 1.42 NA objective, and an ImageEMX2 digital camera or Hamamatsu Orca Fusion CMOS digital camera. Images were acquired with Metamorph software version 7.10.3279 (Molecular Devices).

### Cell culture

All cells were maintained at 37°C and supplemented with 5% CO2. HeLa human cervical adenocarcinoma originally from the American Type Culture Collection (ATCC; Manassas, VA, USA) were grown in Dulbecco’s modified Eagle’s medium (DMEM) (no. 11965-084; Gibco, Carlsbad, CA, USA) supplemented with 9% v/v fetal bovine serum (FBS; Atlanta Biologicals, S11150).

### DNA transfection

Cells were suspended by trypsinization and transfected using Lipofectamine 2000 (Invitrogen, no. 11668027) according to the manufacturer’s instructions.

### Live cell imaging

Cells were seeded in a four-chamber, no. 1.5 glass-bottom dish (Cellvis) and cultured as appropriate for the intended experiment. Before imaging, the medium was replaced with a warm imaging solution [DMEM, high glucose, HEPES, no phenol red (Gibco, 21063029) supplemented with 10% v/v fetal bovine serum]. The dish was immediately transferred to a Tokai Hit (model: STXG-WSKMX-SET) stage-top incubator preheated to 37°C. The dish was allowed to equilibrate for 10 min before initiating acquisition.

### Immunofluorescence

Cells were fixed in 4% w/v paraformaldehyde in phosphate-buffered saline (PBS) for 15 min at room temperature, rinsed with PBS, and permeabilized in 0.1% v/v Triton X-100 (Pierce Biotechnology). Cells were immunolabeled in blocking solution for 1 hour at RT with antibody against Sec31A (Transduction lab, 1:1000). After rinsing with PBS, goat secondary antibodies conjugated to Alexa Fluor 488 fluorescent dyes (Molecular Probes, Carlsbad, CA, USA) were diluted to 1 μg/ml in blocking solution and were added to cells for 45 min.

### Image Analysis

Custom-made scripts were used to perform puncta recognition analysis in Python. We started by using Machine learning to train our model (Yolov5 developed by Ultralytics (https://github.com/ultralytics/yolov5.git) for GUV recognition. Next, we used this trained model to recognize individual GUVs from confocal image frames with multiple GUVs. We used the corresponding protein channels for puncta recognition for each recognized GUV. In each protein channel, we performed background subtraction based on the background intensity determined using minimum cross entropy thresholding (threshold_li from scikit-image). Next, we performed Gaussian noise reduction on the background-subtracted images using the Fast Nl Means Denoising algorithm followed by Gaussian Blur to get a regularly shaped punctum. Subsequently, we adjusted the image contrast to sparse out the image pixel values (using Equalize Bar chart from OpenCV) before generating a binary mask (using threshold_otsu from scikit-image, which minimizes the intra-group pixel value variance). In the foreground pixels, we counted groups of connected pixels that are larger than 5 pixels as individual punctum. We calculated the proportion of GUVs with puncta as the ratio of GUVs that have at least one punctum with the total number of recognized GUVs. Finally, for each recognized punctum, we generated a rectangular bounding box. We considered a punctum as colocalized among different protein channels if their bounding boxes overlapped. The proportion of colocalized puncta was calculated as the colocalized punctum divided by the total puncta in the channel with the highest puncta.

The representative images in Figures 1B and 5 were background subtracted before contrast adjustment for clear puncta depiction. Specifically, the representative figures were thresholded to remove the background fluorescence from the unbound fluorescent protein and depict the puncta observed on the periphery of GUVs. For each representative image, a square area of approximately 10 pixels was selected outside the periphery of the GUV in ImageJ. The average pixel value within this selected area was considered the background fluorescence intensity and subtracted from the entire channel. Similar thresholding was done for each (both protein and lipid) channel. Finally, the LUTs for the backgrounded subtracted channels were marginally adjusted to improve the contrast of the puncta on the GUV periphery. This does not affect our quantification analysis which has been described previously.

All image analyses for cell experiments were performed in ImageJ. To quantify for mCherry-ALG-2 positive area, a threshold was empirically determined for individual cells with different expression levels. ImageJ command *Analyze Particle* was then used to measure above threshold particle area per cell.

### Molecular dynamics simulations

Molecular dynamics simulations were performed with GROMACS 2020 (33) using the CHARMM36m force field (34). Atomistic models of the apo and Ca^2+^-bound forms of ALG-2 were based on crystal structures with PDB IDs 2ZND and 2ZN9 (15), respectively. Each subunit of the dimeric models consisted of residues 21-191. The N-terminal Gly/Pro-rich region, shown to be dispensable for ALG-2 membrane binding in GUV-based experiments was excluded from our structural models. The exposed amino group of Ala21 in the truncated construct was neutralized. Mutant structures were generated in PyMOL (35). Protonation states of amino acid side chains were assigned according to pKa prediction by PROPKA (36). Six 14 x 14 nm^2^ patches of membrane each consisting of a random distribution of 70% DOPC and 30% DOPS coarse-grained lipids were independently prepared using the *insane* method (37), solvated with 150 mM of aqueous NaCl, equilibrated for 200 ns, and converted into an atomistic representation using the CG2AT2 tool (38). Atomistic structures of wild-type or mutant ALG-2 were placed above the resulting membranes with a minimum distance of ∼2 nm. Each protein-membrane system was subsequently subjected to re-solvation and 10 ns of further equilibration. During equilibration, harmonic positional restraints with a force constant of 1000 kJ mol^-1^ were applied to non-hydrogen protein atoms or backbone beads. For the Ca^2+^-bound form of wild-type or mutant ALG-2, harmonic restraints with a force constant of 100 kJ mol^-1^ were additionally applied to the distance between each calcium ion and the center of mass of its coordinating atoms during simulations. The system temperature and pressure were maintained at 310 K and 1 bar, respectively, using the velocity-rescaling thermostat (39) and a semi-isotropic Parrinello-Rahman barostat (40) in the production phase. The integration time step was 2 fs. Long-range electrostatic interactions were treated using the smooth particle mesh Ewald method (41, 42) with a Fourier spacing of 0.12 nm and charge interpolation through fourth-order B-splines. We applied a real-space distance cut-off of 1 nm to non-bonded interactions. The LINCS algorithm was used to constrain covalent bonds involving hydrogen atoms (43). Simulation trajectories were analyzed through MDAnalysis 2.0 (44, 45) in Python 3.6.

### Poisson-Boltzmann calculations

Electrostatic potential maps were calculated for apo and Ca^2+^-bound ALG-2 by numerically solving the linearized Poisson-Boltzmann equation (46) using the Adaptive Poisson-Boltzmann Solver (47, 48). The multigrid finite-difference calculation (49) was performed on a 129 x 129 x 129 grid with dimensions 13 x 13 x 13 nm^3^, applying Debye-Hückel boundary conditions, and the finer grid focused into 10 x 10 x 10 nm^3^. Protein atoms were assigned radii and partial charges from the CHARMM36m force field (50). The radius of an implicit solvent molecule was set to 0.14 nm, the ionic strength to that of 0.15 M NaCl, and the dielectric constant to 78.5 for the solvent and 2 for the protein.

### Modeling analysis

AlphaFold2 (https://alphafold.ebi.ac.uk/) was run using the ColabFold notebook (https://colab.research.google.com/github/sokrypton/ColabFold) using version v1.5.2 on default settings. ChimeraX 1.3 was used to overlay the AlphaFold predicted R34E/K37E/R39E ALG-2 with des3-23 WT ALG-2 (PDB – 2ZN9) as shown in Fig. S1.

### Statistical analysis

Statistical analysis was performed with GraphPad Prism 9.0 (La Jolla, CA, USA). The data of the GUV binding assay was analyzed by student’s t-test and one-way ANOVA. The significance between the two calculated areas was determined using a student’s two-tailed unpaired t-test. *P* < 0.05 was considered statistically significant.

## Acknowledgments

We thank Liv Jensen for help with data analysis script.

## Supplementary Information for

**This PDF file includes**: Figure S1 to S5

**Fig. S1.**
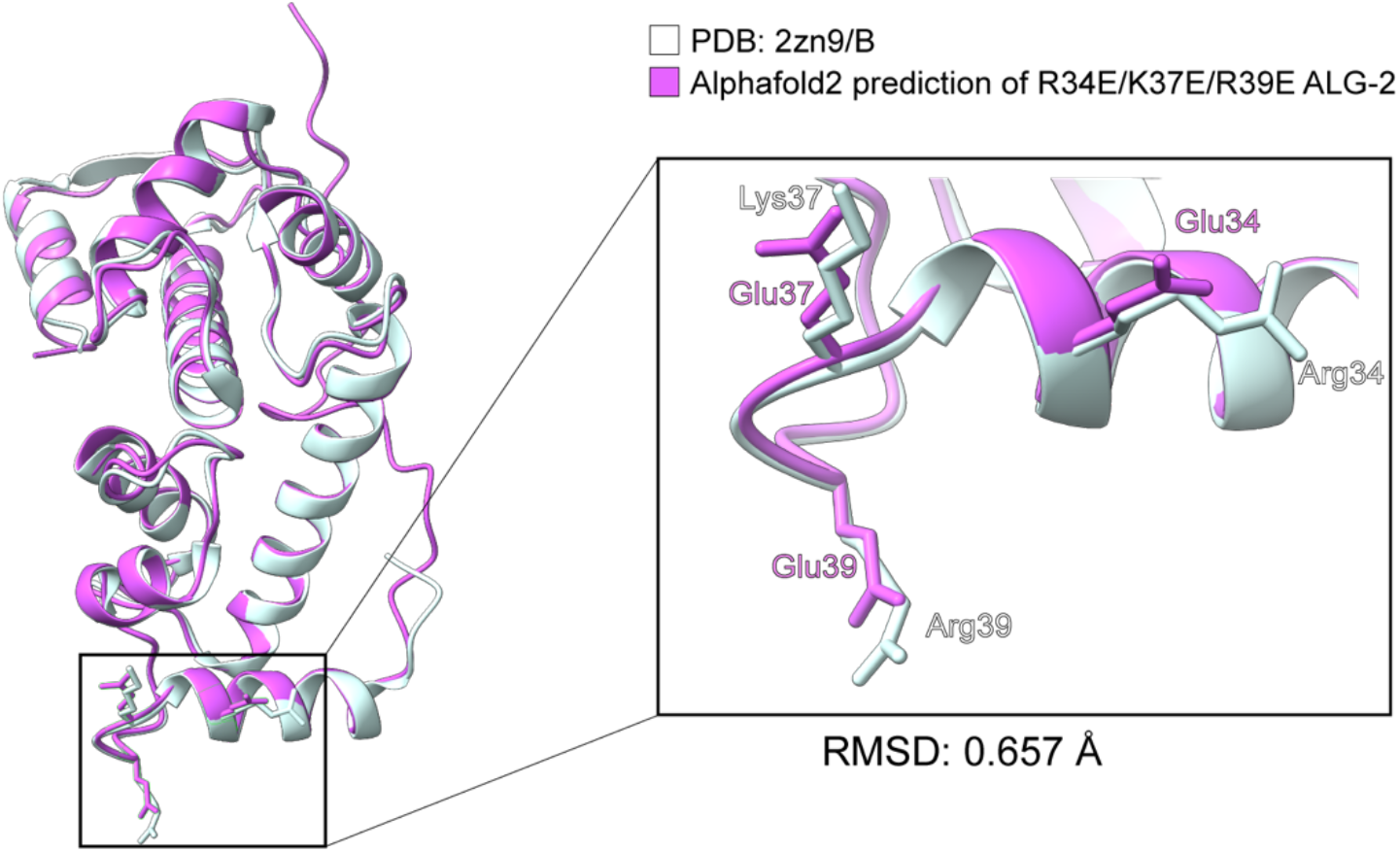
AlphaFold comparison between WT and R34E/K37E/R39E ALG-2 mutant. Overlay of the crystal structure of des3-20 WT ALG-2 (PDB – 2zn9: B) (white) with AlphaFold2 (AF2)–predicted model for R34E/K37E/R39E ALG-2 (magenta). The C_α_ comparison between residues 24 to 188 (which are ordered) between the des3-20 WT ALG-2 (PDB – 2zn9: B) and AF2–predicted model R34E/K37E/R39E ALG-2 mutant, resulted in a RMSD value of 0.657Å and a TM-Align score of 0.978 (when normalized by length of des3-20 WT ALG-2 (PDB – 2zn9: B)). RMSD, root mean square deviation. TM-score, template modeling score.

**Fig. S2.**
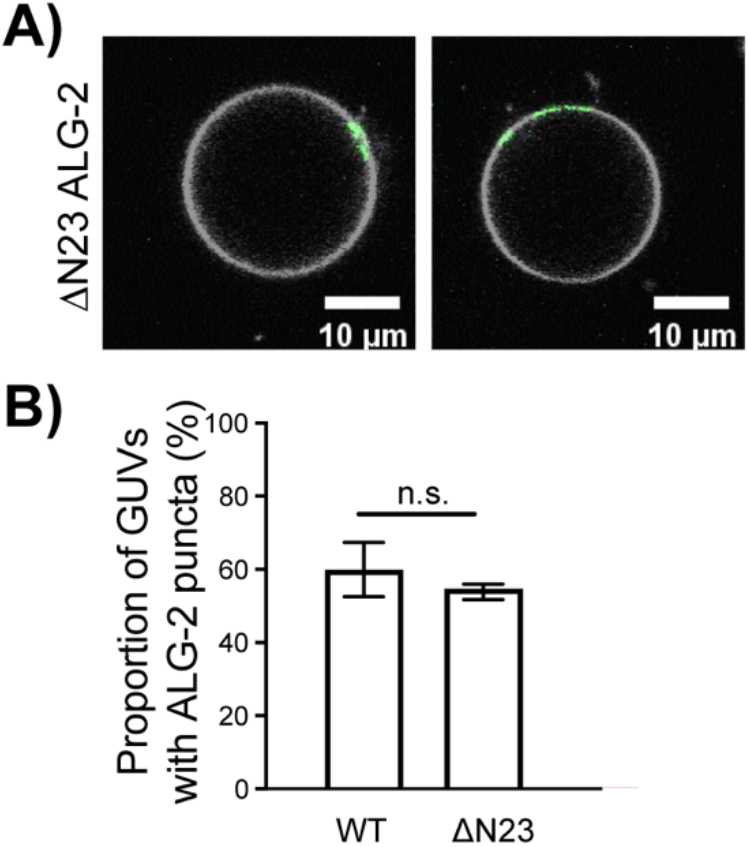
Membrane binding of N-terminally deleted ALG-2. The ΔN23 ALG-2 fluorescently labeled with Atto 488 was incubated with 30% DOPS containing GUVs. (A) ΔN23 ALG-2 (green) was recruited to the 30% DOPS GUVs (white). The images are depicted as a merged channels between the ALG-2 and membrane channel. (B) The proportion of GUVs that had at least one ΔN23 ALG-2 punctum (green) on their periphery (white) were plotted for fluorescently labeled WT (n = 1388 GUVs) and ΔN23 ALG-2 (n = 603 GUVs). The data are shown as mean ± SD (vertical line). All results are from at least three independent experiments.

**Fig. S3.**
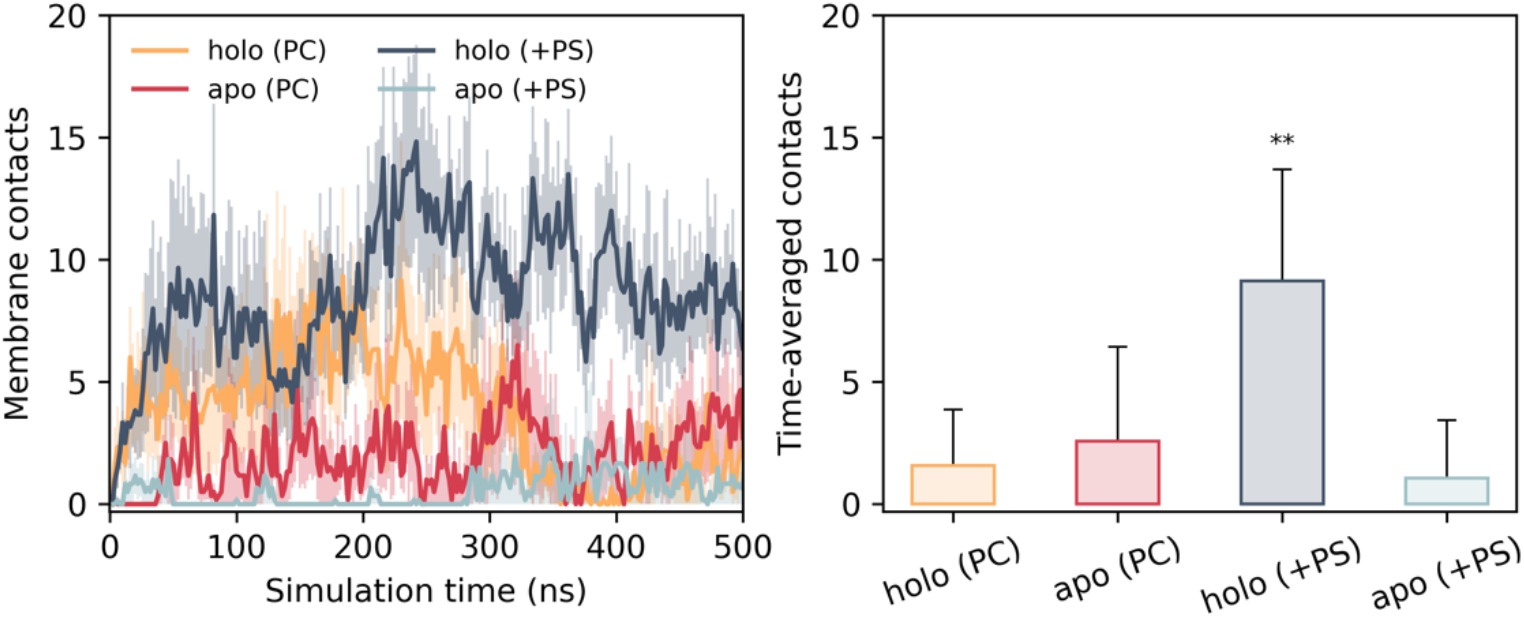
Molecular dynamics simulations of WT ALG-2 with PC only and PC/PS membrane. Number of ALG-2 residues forming membrane contacts during all-atom molecular dynamics simulation replicates, comparing between the Ca^2+^-bound (holo) and apo forms of ALG-2 binding to PC membranes and to membranes containing 30% PS. The mean (solid lines) and standard errors (semi-transparent shading) are plotted over time for six simulation replicates. Time-averaged membrane contacts and standard deviations are also calculated between *t* = 300 ns and 500 ns of simulation replicates. A statistically significant (0.001 < *p* < 0.01; one-tailed Student’s t-test) difference in binding, compared with the holo protein docking to PC membranes, is denoted by an asterisk.

**Fig. S4.**
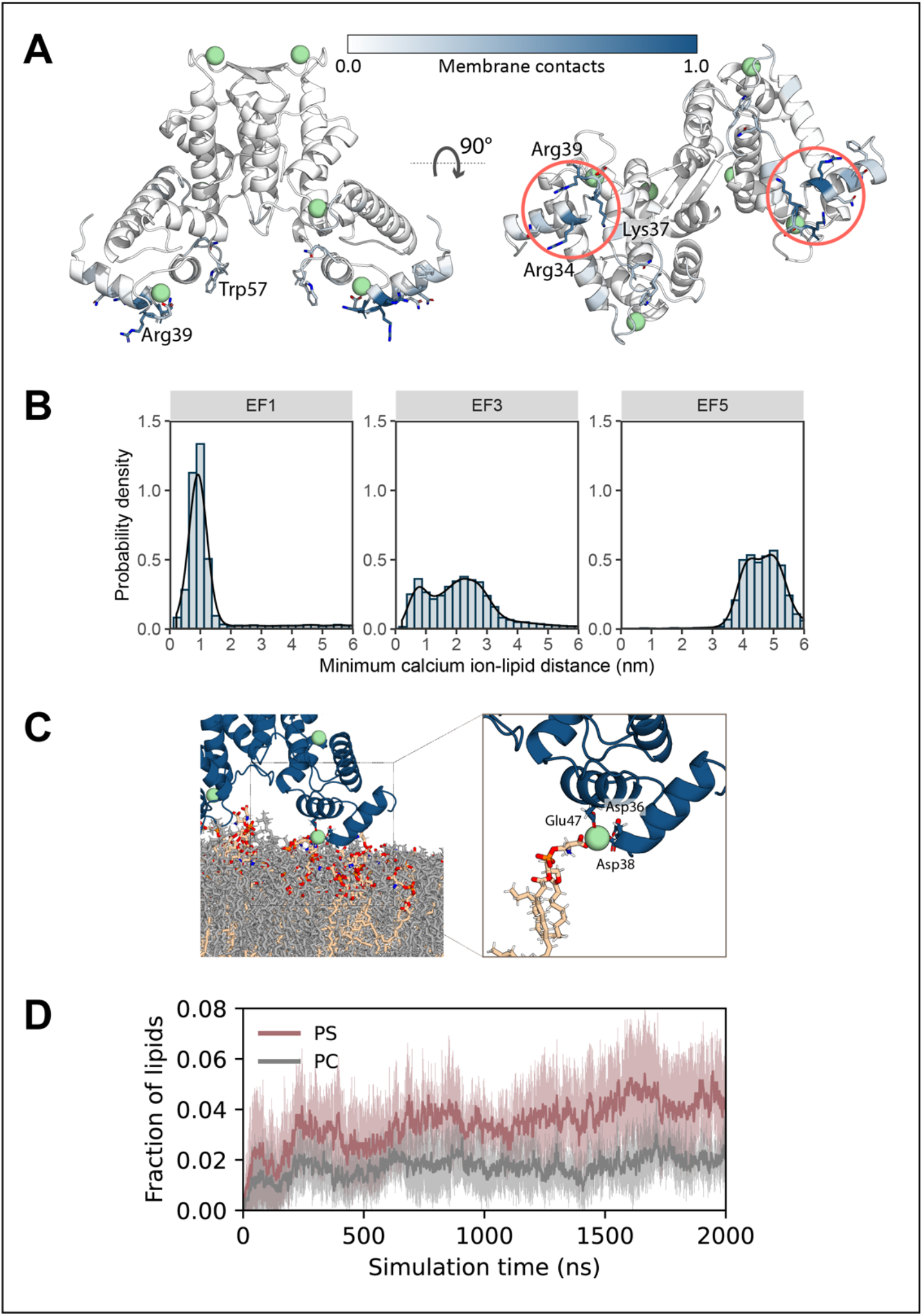
Molecular dynamics simulations of ALG-2 membrane interactions. (A) Structure of Ca^2+^-bound ALG-2 (PDB ID: 2ZN9) with residues colored by their mean frequency of membrane contacts during the final 1 µs of each of six 2 µs simulation replicates (white to blue at increasing contact frequency). (B) Distribution of distances between Ca^2+^ ions bound to EF1, EF3, and EF5 and their nearest lipid atom through six 2 µs simulation replicates. (C) Molecular dynamics simulation snapshot capturing direct Ca^2+^ coordination by phosphate oxygen atoms of a membrane PS lipid. (D) Fractions of the number of membrane PC and PS lipids, respectively, that are in contact with ALG-2 over time. Showing the mean values across six simulation replicates, with the standard deviations indicated as shaded gray and pink bands.

**Fig. S5.**
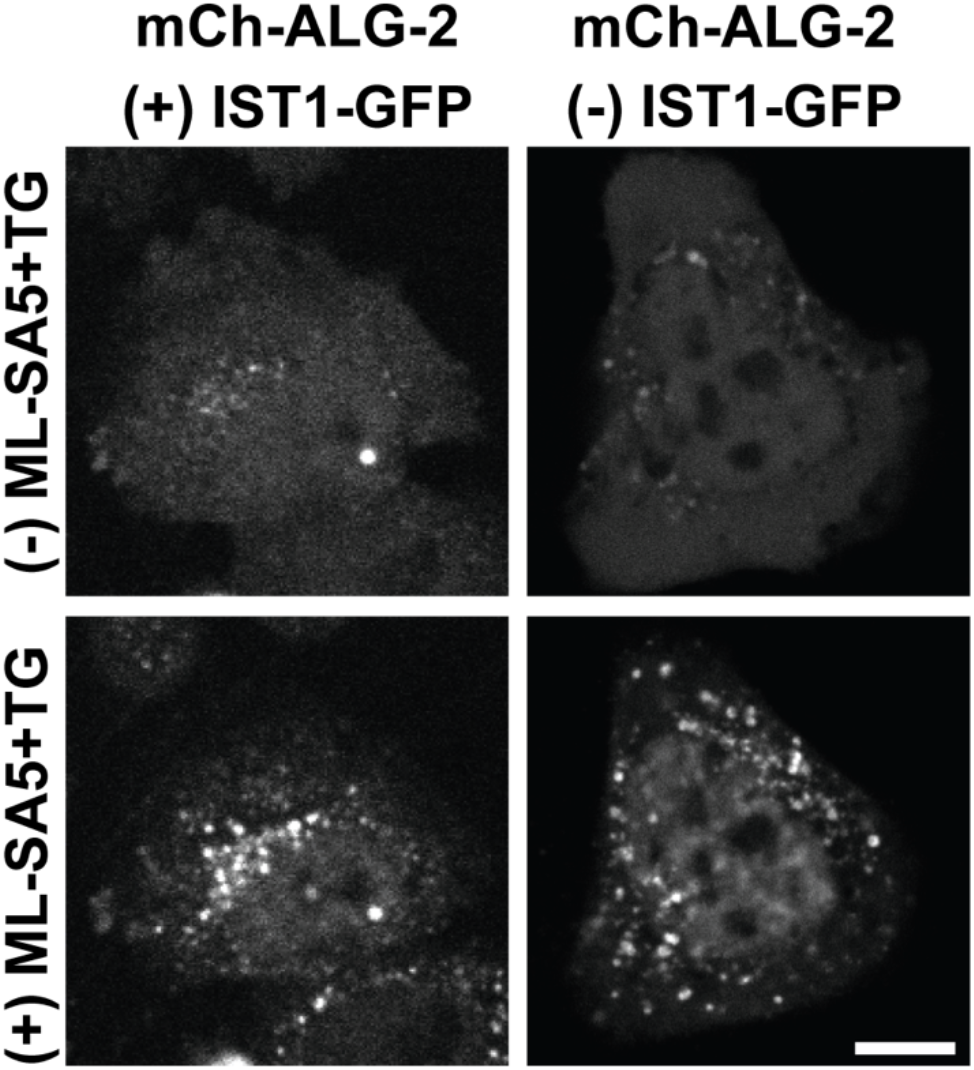
ALG-2 overexpression in the absence of IST overexpression. ALG-2 KO HeLa cells were transfected with mCherry-ALG-2 alone or together with IST1-GFP. Cells were imaged live before and 15min after addition of ML-SA5 and TG. The scale bar is 10 μm.

